# A Single-Cell Peripheral Immune Atlas Spanning High-Risk Lesions to Invasive Breast Cancer in Black and White Women

**DOI:** 10.64898/2026.05.04.722801

**Authors:** Fangyuan Chen, Esther Ritah Ogayo, Tasnim Rahman, Abigail Recko, Hanna Starobinets, Milos Spasic, Adrienne Marie Parsons, Peter van Galen, Elizabeth A. Mittendorf, Sandra S. McAllister

**Affiliations:** Breast Oncology Program, Dana-Farber Cancer Institute, Boston, MA, USA; Hematology Division, Brigham and Women’s Hospital, Boston, MA; Harvard Medical School, Boston, MA, USA; Ludwig Center at Harvard, Harvard Medical School, Boston, MA, USA; Harvard Stem Cell Institute, Cambridge, MA, USA; Broad Institute of Harvard and MIT, Cambridge, MA, USA; Division of Breast Surgery, Department of Surgery, Beth Israel Deaconess Medical Center, MA, USA

## Abstract

Black women are at risk for breast cancer nearly a decade before women of other racial groups for unclear reasons. Because immune responses influence cancer initiation and progression, we performed single-cell RNA sequencing of peripheral blood mononuclear cells from non-Hispanic Black (NHB) and non-Hispanic White (NHW) women with high-risk breast lesions, ductal carcinoma in-situ, and invasive breast cancer. Race-associated transcriptional differences were observed across all disease states and were most pronounced in invasive disease. Computational analyses, supported by flow cytometric protein analysis, revealed enrichment of chronic inflammation, immune regulatory programs, and immune aging pathways in NHB cancer patients, particularly in monocytes, dendritic cells, CD4^+^ T cells, and B cells. From these data, we derived a population-level immune signature (IMM-POP) comprising genes differentially enriched in this subset of immune cells from NHB breast cancer patients. IMM-POP correlates with an immunosuppressive signature in external breast cancer datasets. We thus provide a single-cell peripheral immune atlas integrating race and breast disease state.

**Significance:** This study revealed race-specific peripheral immunity features in precancerous and invasive breast cancers: Black patients exhibited features of chronic inflammation and immune aging compared with White patients, suggesting ‘immune weathering’ and providing insights for studying early onset of breast cancer in Black patients.

## Introduction

Non-Hispanic Black (NHB) women are diagnosed with breast cancer at a younger age(1) and have higher age-adjusted breast cancer mortality(2) than women of other racial groups. Biological factors underlying these differences are likely multifactorial but remain unclear(3). Progress is hindered by a lack of high-resolution data from precancerous and early-stage disease across diverse racial groups, thus limiting insights into when or if biological differences arise.

The immune system plays a central role in cancer control(4). Under homeostatic conditions, immune cells recognize and eliminate pre-malignant cells, preventing tumor initiation(5). However, once cancer develops, tumor cells can evade or suppress anti-tumor immune responses(6,7). Immune characterization in breast cancer is largely derived from analyses of tumor tissue, providing important insights into local tumor-immune interactions(8). Peripheral immune profiling, however, enables assessment of systemic immune features in the high-risk and early disease settings for which tissue is often not obtained or available. Nevertheless, whether peripheral immune profiles vary across early disease states or between populations is not clear.

We asked whether peripheral immune profiles differ between NHB and NHW women across at-risk to early-stage breast cancer, potentially informing biological factors associated with earlier cancer onset experienced by NHB women. We assembled a cohort of NHB and NHW women with high-risk breast lesions (HRL), ductal carcinoma in situ (DCIS), and early-stage breast cancer (BC) and performed single cell RNA- sequencing (scRNA-seq) of peripheral blood mononuclear cells (PBMCs). We identified: (1) transcriptional differences across disease states independent of race; (2) distinct race-associated transcriptional programs in each disease state; and (3) enrichment of inflammatory and immune regulatory programs from specific immune lineages in NHB compared with NHW patients with invasive breast cancer. We derived a population-level gene signature, IMM-POP, that captures immune-related transcriptional features in NHB relative to NHW breast cancer patients and shows significant co-expression with an immunosuppressive signature in independent breast cancer cohorts. Our dataset provides a high-resolution, racially diverse resource for investigating systemic immune variation across early breast disease.

## Results

### Single-cell transcriptomics of peripheral blood mononuclear cells across disease states and racial groups

PBMC samples were curated from age-matched self-identified non-Hispanic Black (NHB; n=27) and non-Hispanic White (NHW; n=28) women with high-risk lesions (HRL), ductal carcinoma in situ (DCIS), or invasive breast cancer (BC) (Fig. 1a,b). The HRL cohort included 20 patients (10 NHB, 10 NHW) enrolled in our institution’s B-PREP program(9) who were at increased risk for breast cancer, defined by biopsy-proven atypical ductal hyperplasia, atypical lobular hyperplasia, or lobular carcinoma in situ (Supplementary Table 1). The DCIS and BC groups included patients enrolled through Project SHARE(10). The DCIS group (4 NHB, 6 NHW) included patients with estrogen receptor–positive (ER+)

**Figure 1.**
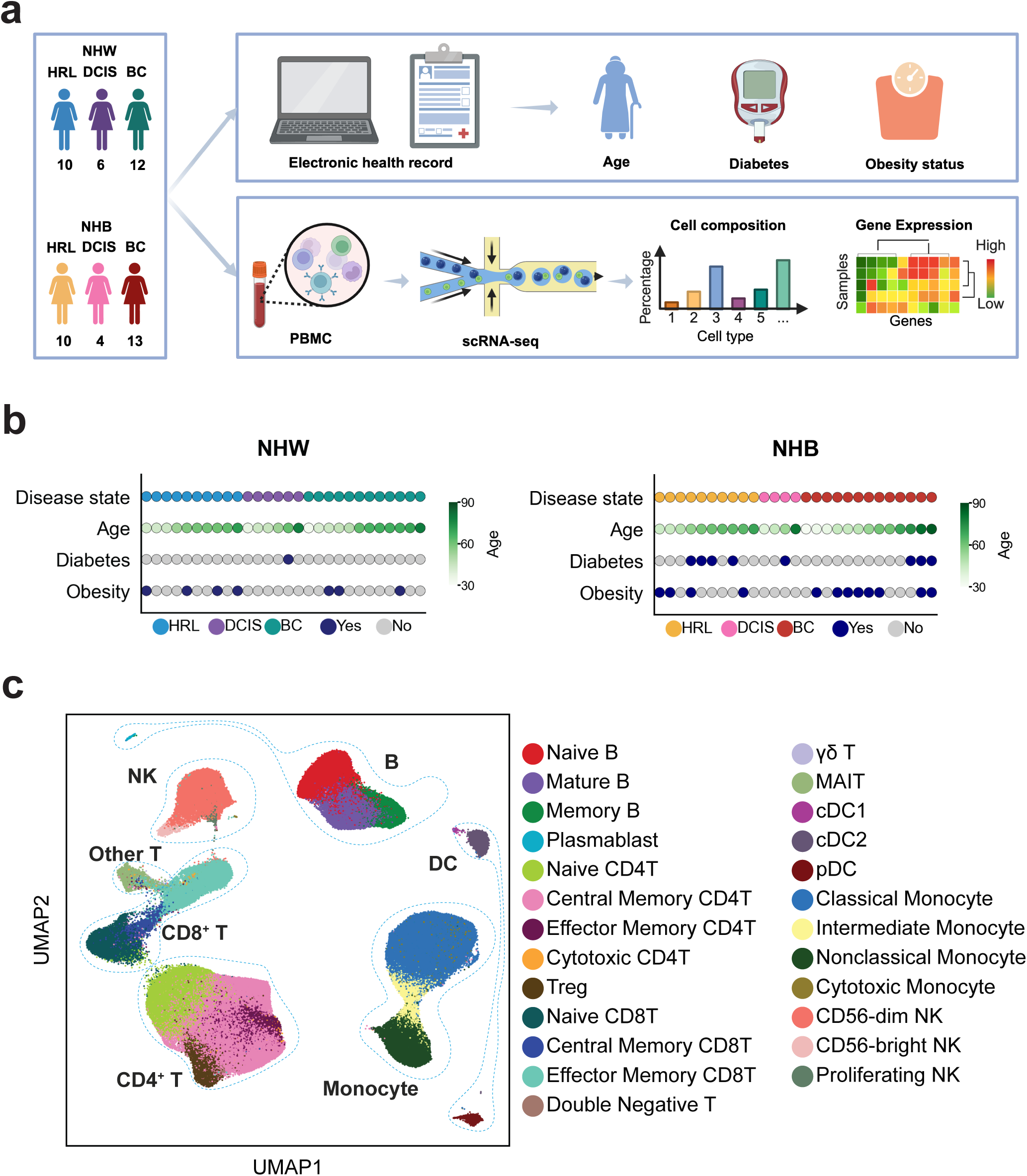
Single-cell transcriptional profiling of PBMCs from NHB and NHW patients across 3 breast disease states. **a**, Overview of patient cohorts and PBMC analysis workflow (created in BioRender; Chen, F. (2025), https://BioRender.com/mvwqlws). **b**, Clinical characteristics of NHW (left panel) and NHB (right panel) patients; dots represent individual patients and are placed in same order for each parameter. **c**, UMAP plot of PBMC cell types from all patients. Blue dashed lines encompass indicated major immune cell types. HRL, high-risk breast lesion; DCIS, ductal carcinoma in situ; BC, breast cancer; NHW, non-Hispanic White; NHB, non-Hispanic Black; PBMC, peripheral blood mononuclear cell; scRNA-seq, single-cell RNA sequencing; UMAP, Uniform Manifold Approximation and Projection; cDC1, conventional type 1 dendritic cell; cDC2, conventional type 2 dendritic cell; pDC, plasmacytoid dendritic cell; NK, natural killer cell; MAIT, mucosal-associated invariant T cell.

DCIS, and the invasive breast cancer group (13 NHB, 12 NHW) consisted of patients with pre-treatment, early-stage ER+ invasive breast cancer (Fig. 1a; Supplementary Table 2). All patients were verified to be free of acute infections, vaccinations, or other concurrent cancers at the time of blood collection, which occurred from shortly after initial diagnosis to within one year thereafter. Age, diabetes, and obesity status were obtained from the electronic medical record at time of blood collection (Supplementary Table 1), and histopathological information of DCIS and BC was obtained from pathological report (Supplementary Table 2). Among comorbid conditions, 6 patients had diabetes alone, 16 had obesity alone, and 3 had both (Fig. 1b, Supplementary Table 1).

We performed single-cell RNA-sequencing (scRNA-seq) on the PBMC samples and generated a transcriptomic dataset consisting of 434,565 cells representing 25 functional minor immune cell types from five major immune cell types: B cells, T cells, monocytes, dendritic cells (DCs), and natural killer (NK) cells (Fig. 1c). Standard quality-control metrics, including per-cell gene counts, unique molecular identifier counts, mitochondrial gene percentage, and major cell type prediction scores (Supplementary Fig. S1a–d), indicated high quality of the dataset. Expression of major immune cell type-specific proteins (computed by Azimuth using the CITE-seq PBMC reference dataset(11)) and marker genes of each cell type (using differential gene expression analysis) were consistent with our cell type annotations (Supplementary Fig. S1e,f).

### PBMC composition differs by both disease status and racial group

The relative contributions of major immune cell types: B cells, CD4⁺ T cells, CD8⁺ T cells, other T cells (including γδT cells, double-negative T cells, and mucosal-associated invariant T [MAIT] cells), monocytes, dendritic cells (DCs), and natural killer (NK) cells as a proportion of total PBMCs were similar across disease states and race (Supplementary Fig. S2a, b), indicating compositional stability among major immune lineages.

Further analysis of 25 immune cell functional subtypes across disease states within each racial group (Fig. 2a) revealed statistically significant differences in abundance (Supplementary Table 3). Among NHW patients, disease state-specific differences were observed across several T cell and NK cell subtypes: compared with the HRL group, patients with DCIS had reduced proportions of naïve CD4^+^ T cells (Fig. 2b), and those with BC had lower proportions of central memory CD4^+^ T cells (Fig. 2c). In parallel, cytotoxic CD4^+^ T cells(12) comprised a smaller proportion of total T cells in BC than in DCIS (Fig. 2d). Also in the NHW patients, breast cancer was associated with increased abundance of mature cytotoxic CD56^dim^ NK cells(13) (Fig. 2e) and decreased abundance of the cytokine producing, immunoregulatory CD56^bright^ NK cells(14) relative to HRL(Fig. 2f).

**Figure 2.**
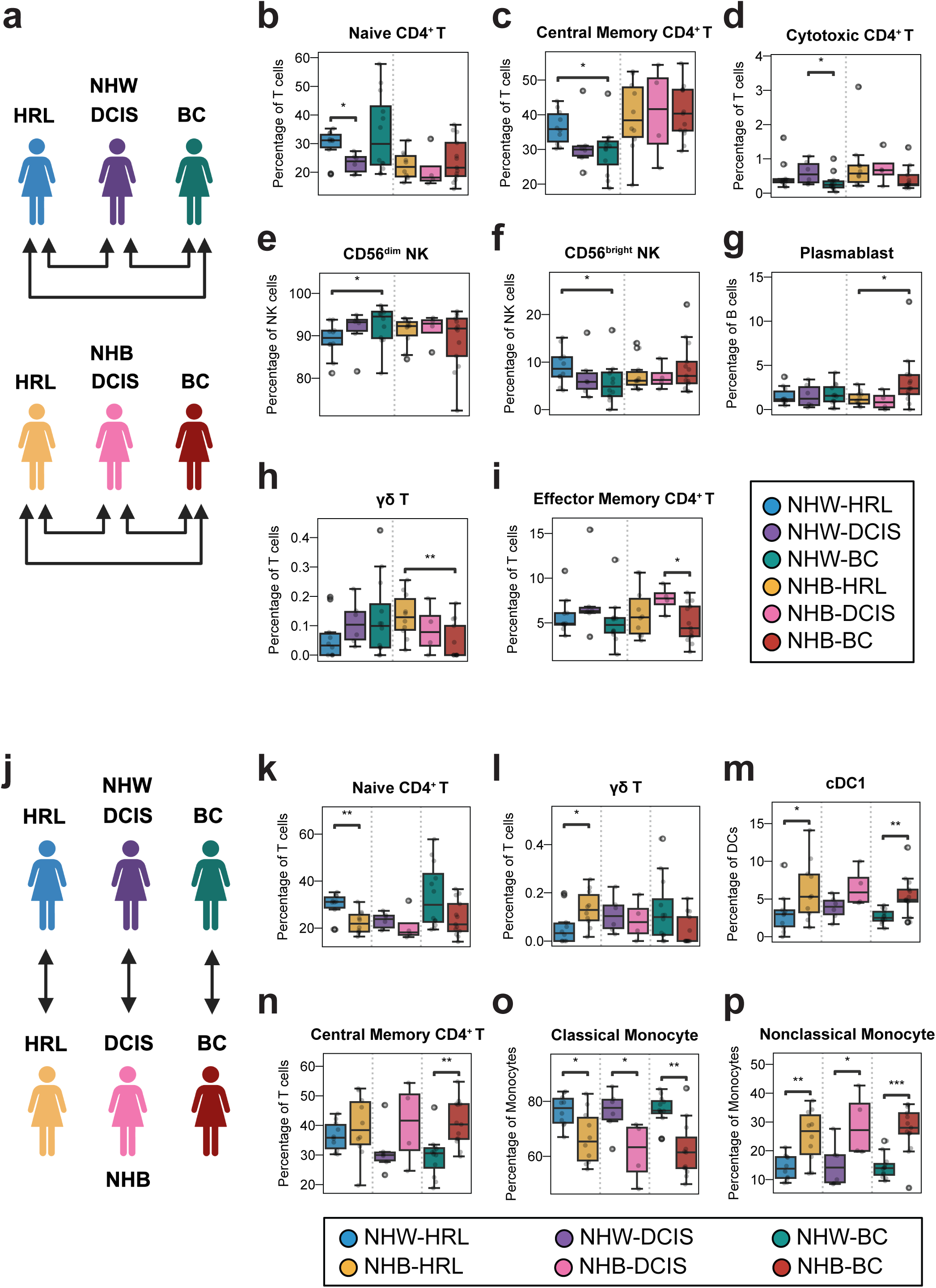
PBMC composition across disease states and racial groups. a, Cohort schematic for comparing PBMCs across disease states within each race group. **b-i**, Percentage of PBMC cell types within the indicated parental immune lineage for comparisons in (a). **j**, Cohort schematic for comparing PBMCs between racial groups within each disease state. **k-p**, Percentage of PBMC cell types within the indicated parental immune lineage for comparisons in (j). Mann-Whitney U test; *p ≤ 0.05, **p ≤ 0.01, ***p ≤ 0.001, ****p ≤ 0.0001. HRL, high-risk breast lesion; DCIS, ductal carcinoma in-situ; BC, breast cancer. BioRender. Chen, F. (2026) https://BioRender.com/gyb1uoo.

The NHB patients exhibited a distinct pattern of disease-associated immune remodeling. Relative to HRL, the BC group had a higher proportion of plasmablasts (Fig. 2g), a short-lived, terminally differentiated, and antibody-secreting B cell subtype(15), and reduced proportions of γδ T cells (Fig. 2h), which perform innate-like immune surveillance(16). Also in the NHB patients, effector memory CD4^+^ T cells(17) were less abundant in breast cancer than in DCIS (Fig. 2i). No other immune cell subpopulations differed significantly in abundance across disease states in either racial group (Supplementary Fig. S2c).

In addition to the comparisons across disease states within each race group above, we compared immune cell abundance between NHB and NHW patients within each distinct disease state (Fig. 2j). Within the HRL group, NHB patients had lower proportions of naïve CD4^+^ T cells (Fig. 2k) and higher proportions of γδ T cells (Fig. 2l). Type 1 conventional dendritic cells (cDC1), which mediate antigen cross-presentation and cytotoxic T-cell priming(18), were more abundant in NHB than in NHW patients within both the HRL and BC groups (Fig. 2m). In BC, central memory CD4^+^ T cells were more abundant in NHB than in NHW patients (Fig. 2n). Notably, across all disease states, classical monocytes, key mediators of phagocytosis and innate inflammatory responses(19), constituted lower proportions and non-classical monocytes, which are involved in vascular homeostasis, immune surveillance, and persistent inflammation(20), were more abundant in the NHB patients (Fig. 2o, p). No significant race-associated differences were detected in the remaining immune cell subtypes (Supplementary Fig. S2d).

Together, these results demonstrate that although cell counts of major immune lineages remain stable across racial groups and disease states, selective PBMC functional subpopulations differ in proportion by disease state and by race, underscoring the value of high-resolution peripheral immune profiling for detecting immune variation not evident at the lineage level.

### Disease state-associated transcriptional remodeling of peripheral immune cells within racial groups

To quantify transcriptional differences across disease states within each racial group, we aggregated the number of DEGs in each cell type for pairwise comparisons (DCIS vs. HRL, BC vs. DCIS, and BC vs. HRL) (Fig. 2a). NHB patients showed greater transcriptional differences between BC and either HRL or DCIS than between DCIS and HRL. Specifically, NHB patients had 6,780 DEGs for BC vs. HRL and 1,342 DEGs for BC vs. DCIS, compared with only 14 DEGs for DCIS vs. HRL (Supplementary Table 4). In the NHW patients, there were 2,209 DEGs for BC vs. HRL, and only 3 DEGs for BC vs. DCIS; and 4 DEGs for DCIS vs. HRL (Supplementary Table 4).

The pronounced transcriptional changes observed in the breast cancer state, particularly in NHB patients, prompted us to investigate which immune cell types accounted for these variations. We therefore ranked cells by their cumulative number of DEGs across each pairwise disease-state comparison. The top five cell types included classical monocytes (3,315 DEGs), central memory CD4⁺ T cells (3,071 DEGs), non-classical monocytes (2,277 DEGs), naïve CD4⁺ T cells (1,954 DEGs), and effector memory CD8^+^ T cells (1,343 DEGs) (Fig. 3a).

**Figure 3.**
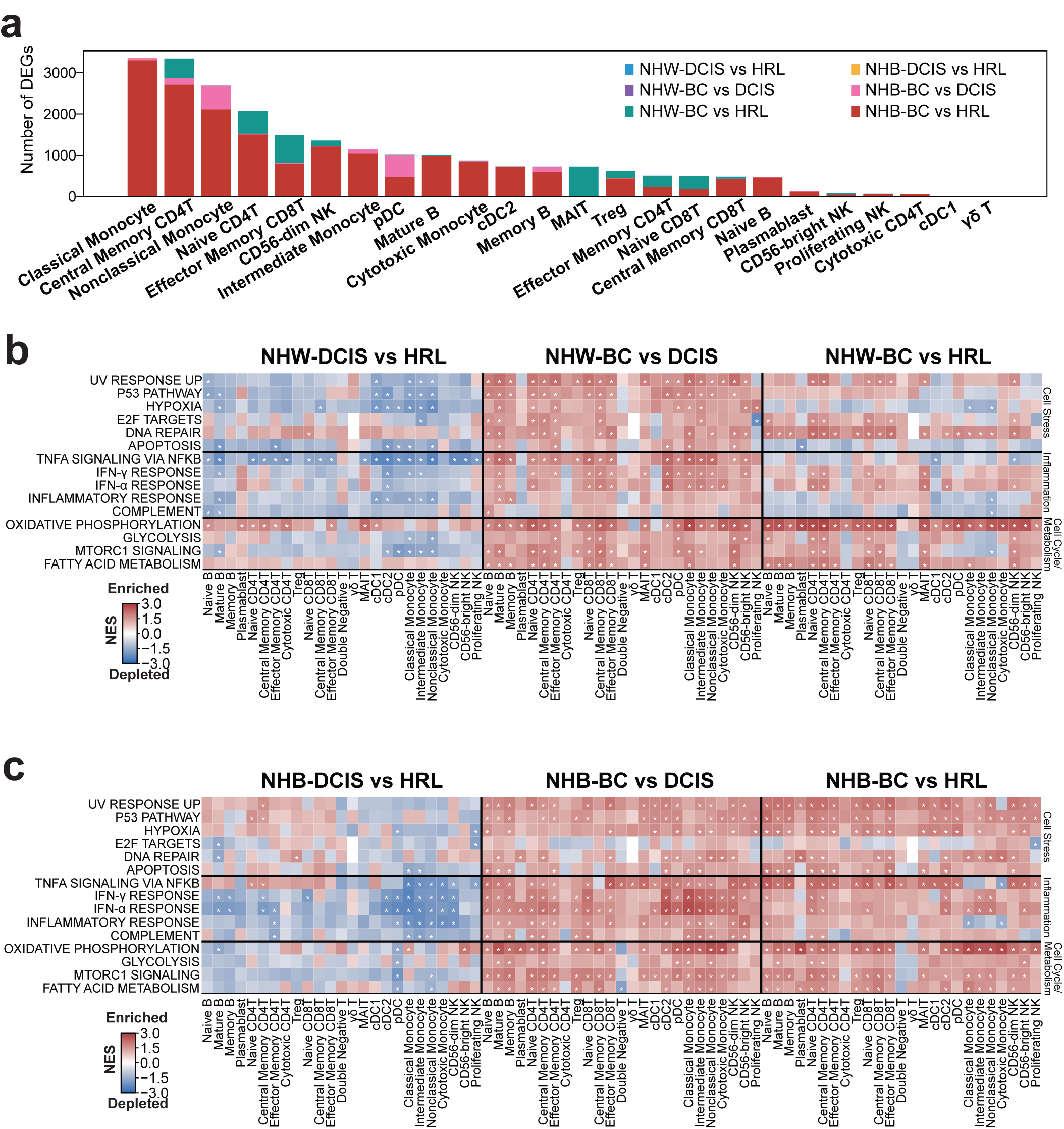
Transcriptional differences in peripheral blood immune cells across disease states in each racial group. **a,** Number of differentially expressed genes in each peripheral blood cell type compared between indicated disease states in NHW and NHB patients. **b-c**, Gene set enrichment analysis of the Molecular Signatures Database (MSigDB) hallmark pathways in each cell type from comparisons between disease states (DCIS vs. HRL, BC vs. DCIS, and BC vs. HRL) in NHW (b) and NHB patients (c). Selected pathways related to cell stress, inflammation, and cell cycle/ metabolism are shown. Box color represents normalized enrichment score (NES); asterisks indicate GSEA FDR-adjusted p value < 0.05.

We also ranked the cell types that accounted for the greatest number of DEGs within each disease state comparison by race. Comparing BC to HRL in the NHB cohort, the cell types with the greatest number of DEGs included classical monocytes (3,299 DEGs), central memory CD4 T cells (2,706 DEGs), non-classical monocytes (2,111 DEGs), naïve CD4^+^ T cells (1,498 DEGs), and CD56^dim^ NK cells (1,209 DEGs) (Fig. 3a, red bars; Supplementary Table 4). Comparing BC to HRL in the NHW cohort, the leading populations included mucosal-associated invariant T (MAIT) cells (718 DEGs), effector memory CD8^+^ T cells (680 DEGs), naïve CD4^+^ T cells (553 DEGs), central memory CD4^+^ T cells (466 DEGs), and naïve CD8^+^ T cells (305 DEGs) (Fig. 3a, green bars; Supplementary Table 4). Notable cell types with the highest number of DEGs between BC and DCIS in the NHB cohort included plasmacytoid dendritic cells (pDC), nonclassical monocytes, central memory CD4 T cells, and memory B cells (Fig. 3a, pink bars; Supplementary Table 4).

Taken together, these data revealed disease state-associated cell types that differed in both abundance and transcriptional state (naïve CD4⁺ T cells, central memory CD4⁺ T cells, and CD56^dim^ NK cells; Fig. 2b,c,e), and others that differed only in disease state-associated transcriptional profile (monocyte subtypes; Supplementary Fig. S2c).

To identify functional pathways that might define cell-specific, disease state-associated transcriptional differences within each race group, we performed Gene Set Enrichment Analysis (GSEA; MSigDB Hallmark pathways(21)). Pathways associated with cell stress, inflammation, and cell cycle/metabolism(8) were enriched in breast cancer relative to HRL or DCIS for both races (Fig. 3b,c; Supplementary Fig. S3a,b). More specifically, in both race cohorts, TNFα–NFκB signaling, IFN-γ response, IFN-α response, and inflammatory response pathways were broadly enriched across lymphoid and myeloid cell lineages in BC compared with DCIS (Fig. 3b,c). For the NHB cohort, these same pathways were significantly depleted in DCs, monocytes and NK subtypes when comparing DCIS to HRL. (Fig. 3c). In contrast, the two racial groups showed distinct enrichment patterns of some cell stress pathways when comparing BC to HRL; in NHB patients, UV response, p53, and hypoxia pathways were enriched across most PBMC cell types; whereas in NHW patients, this pattern was much less pronounced, with only the UV response pathway enriched in a few T cell and NK subtypes (Fig. 3b,c).

Collectively, these analyses revealed enrichment of cell stress, inflammation, and cell cycle/metabolism-related pathways, broadly spanning B cells, T cells, dendritic cells, monocytes, and NK cells in BC relative to pre-cancerous states in both race groups. However, disease state-related transcriptional differences also manifested between the race cohorts, whereby NHB patients showed stronger enrichment of cell stress-associated pathways between BC and HRL compared to NHW patients.

### Race-associated transcriptional differences in peripheral immune cells across disease states

Because analyses across disease states within each race cohort (Fig. 2a) revealed differences between NHB and NHW patients, we next compared cell-specific transcriptional profiles between races within each disease state (Fig. 2j). By first aggregating the total number of DEGs between NHB and NHW patients in each disease state, we observed 1,447 DEGs in breast cancer, 82 in DCIS, and 280 in HRL (Supplementary Table 5). In breast cancer, naïve and mature B cells, classical monocytes, and naïve and central memory CD4^+^ T cells showed the highest numbers of DEGs (Fig. 4a, green bars). In the patients with DCIS, the highest contributions arose from mature B cells and pDCs (Fig. 4a, purple bars). In patients with HRL, DEG enrichment was most prominent in monocyte populations, including classical, intermediate, and non-classical monocytes (Fig. 4a, blue bars). Thus, racial differences in gene expression were dominated by B cells in breast cancer, monocytes in HRL, and relatively limited changes observed in DCIS. Importantly, these transcriptional differences did not consistently mirror shifts in cell abundance. For example, classical monocytes, naïve CD4^+^ T cells, and central memory CD4^+^ T cells exhibited race- and disease-state differences in abundance (Fig. 2o,k,n), whereas naïve and mature B cells showed race-associated transcriptional differences without corresponding changes in abundance (Supplementary Fig. S2d).

**Figure 4.**
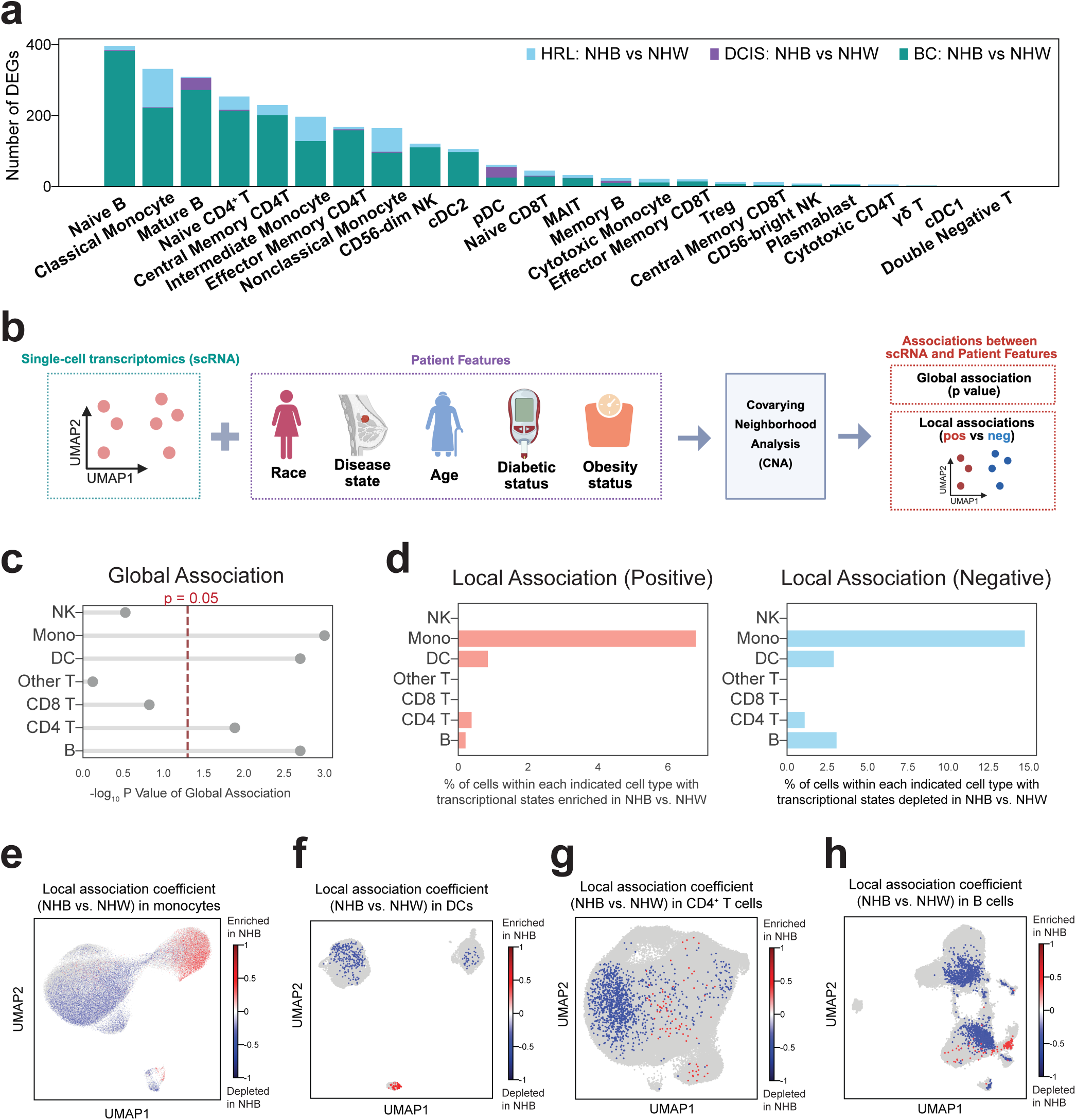
Immune cell transcriptional states that associate with race. **a,** Number of differentially expressed genes in each cell type between NHW and NHB patients in indicated disease states. **b**, Workflow for calculating associations between PBMC transcriptomics with indicated patient features using co-varying neighborhood analysis (CNA) to generate global (one single p value) and local (cell-level FDR-adjusted p value and association coefficient) association metrics. **c**, Global associations between race (NHB and NHW) and transcriptional profiles of each indicated immune cell type (Other T cells include γδT cells, double-negative T cells, and mucosal-associated invariant T (MAIT) cells); −log-(p) values with p = 0.05 (dashed red line) considered significant. **d**, Percentage of indicated immune cells having a transcriptional state associated with race; salmon color (left) = positive association with NHB, sky blue (right) = negative association with NHB. FDR-adjusted p value < 0.1. **e-h**, UMAP plots of the single-cell transcriptomic data colored by CNA local association coefficients in monocytes (e), dendritic cells (f), CD4^+^ T cells (g), and B cells (h). Red dots = individual cells with transcriptional state positively associated with NHB; blue dots = individual cells with transcriptional state negatively associated with NHB (FDR < 0.1); grey dots = cells without significant race associations (FDR ≥ 0.1). BioRender. Chen, F. (2025) https://BioRender.com/kx98ci7.

To complement these approaches, we applied an independent statistical framework, co-varying neighborhood analysis (CNA)(22), which tests whether, and to what extent, transcriptional states of all cells within each major immune lineage are distinguishable by race while adjusting for patient-level covariates. CNA generates two complementary metrics. First, it provides a global association P value, where P < 0.05 indicates an overall race-associated transcriptional difference within a given cell type. Second, it produces local association metrics for individual cells, identifying specific transcriptional states that are disproportionally distributed by race (FDR-adjusted P < 0.1) and indicating whether those states are enriched (positive coefficient) or depleted (negative coefficient) in NHB relative to NHW patients (Fig. 4b). We thus applied CNA to test race-associated transcriptional differences within each major immune lineage, while adjusting for patient-level covariates including age, disease state, diabetes, and obesity, which are known to influence immunity. For each immune lineage, the input included RNA expression in all cells from all 55 patients.

Among the cell lineages, monocytes, DCs, CD4^+^ T cells, and B cells showed both global and local race associations of statistical significance (Fig. 4c-d). Assigning local coefficients to each cell and projecting them onto UMAP for each of these four major lineages revealed distinct race-associated cell state distributions (Fig. 4e-h). Overlaying these coefficients with functional subtype annotations (Supplementary Fig. S4a-d) revealed that cells with transcriptional states enriched in NHB patients mapped primarily to non-classical monocytes, cDC1 cells, and central memory CD4^+^ T cells (Supplementary Fig. S4e–h). In contrast, cells with transcriptional states reduced in NHB patients mapped to classical monocytes, cDC2 and pDCs, and naïve CD4^+^ T cells, as well as an even distribution among mature and naïve B cells (Supplementary Fig. S4e–h).

### Race-associated transcriptional programs across immune cell lineages and disease states

Because race-associated transcriptional states were concentrated in monocytes, dendritic cells, CD4^+^ T cells, and B cells, we sought to identify the functional programs underlying these differences within these lineages. Therefore, we correlated CNA-derived association coefficients with gene expression and performed ImmPort(23) pathway enrichment analysis within each disease state.

Relative to NHW patients, monocytes from NHB patients showed enrichment of inflammatory and immune activation pathways, including interferon signaling, cytokine-mediated responses, Fc receptor-dependent signaling, and antigen presentation pathways, with reduced anti-microbial effector programs across all disease states (Fig. 5a–c). In the HRL and DCIS cohorts, monocytes from NHB patients additionally exhibited reduced reactive oxygen species production, complement activity, and chemotaxis and migration (Fig. 5a, b), suggesting an activated but functionally skewed state in which inflammatory signaling is elevated despite diminished effector and trafficking capacity. In the BC cohorts, monocytes from NHB patients retained enhanced activation (e.g., interferon and Fc receptor signaling) but also showed enriched immunoregulatory pathways, including increased immune cell cross-talk and negative regulation of immune responses (Fig. 5c).

**Figure 5.**
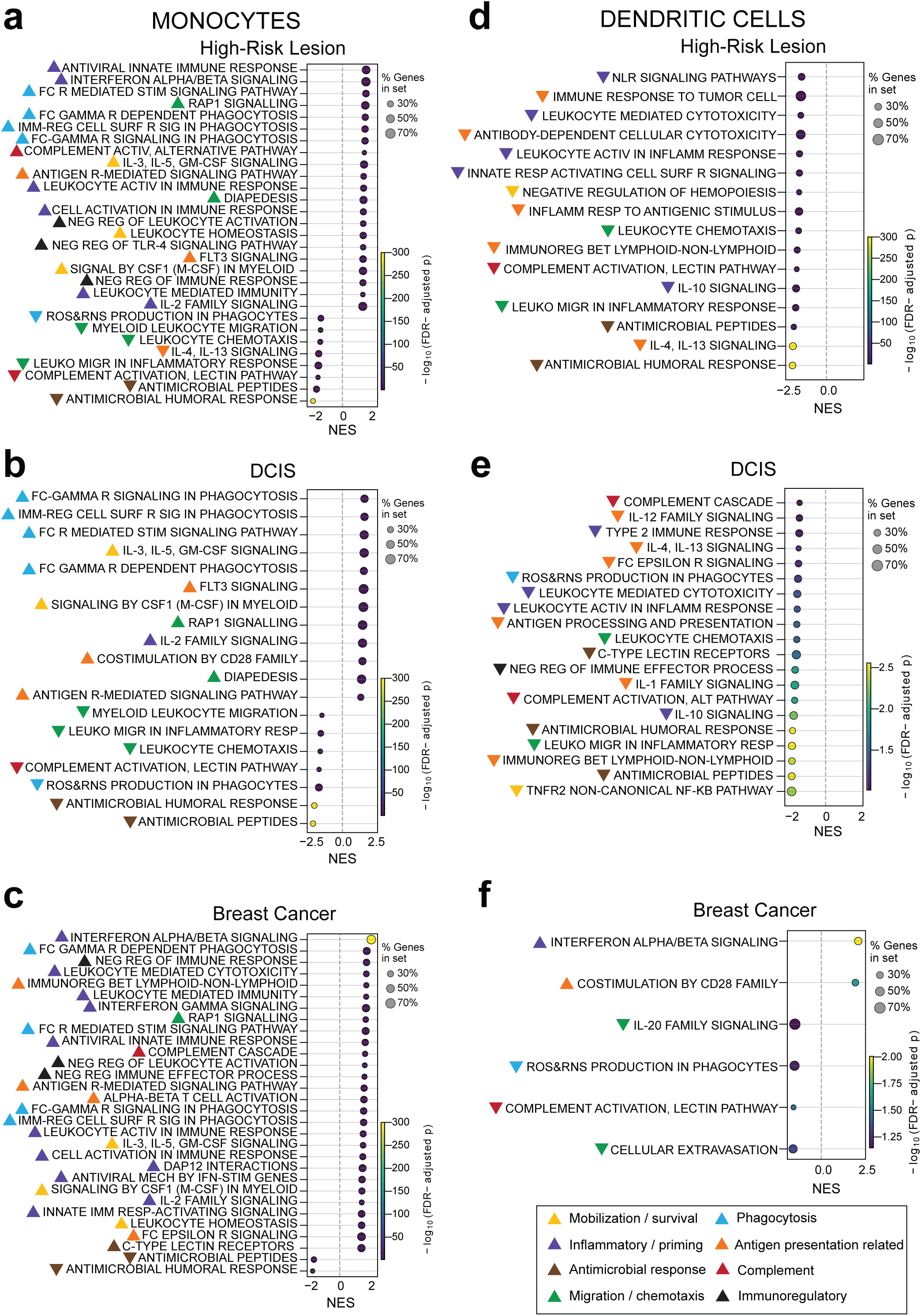
Race-associated pathways across disease states in monocytes and dendritic cells. **a-f**, Pathways enriched (positive NES) and depleted (negative NES) in monocytes (a-c) and DCs (d-f) from NHB relative to NHW patients with HRL (a, d), DCIS (b, e), and BC (c, f) using GSEA and ImmPort databases. Bubble size, percentage of leading-edge genes (genes driving the enrichment signal) in the pathway; bubble color, - log_10_ FDR-adjusted P value; NES, normalized enrichment score. Colored arrows indicate pathways comprising general immune functional categories.

Across disease states, circulating DCs from NHB patients exhibited reduced expression of pathways involved in innate immune sensing, antigen processing and presentation, cytokine signaling, complement activation, antimicrobial responses, and chemotaxis and migration, with no pathway enrichment in the HRL or DCIS groups (Fig. 5d–f). This widespread reduction indicates a broadly attenuated functional profile affecting key processes required for pathogen sensing, immune activation, and cellular trafficking in the NHB patients (Fig. 5d, e). In the BC cohort, DCs from NHB patients additionally displayed enrichment of type I interferon signaling and CD28-mediated costimulatory pathways, suggesting immune priming in the context of diminished innate effector and migratory capacity (Fig. 5f). These observations indicate that circulating DCs from NHB patients share a common pattern of reduced innate and trafficking functions across disease states, with a distinct emergence of priming and T cell-interactive features in invasive BC relative to NHW patients.

Relative to NHW patients, circulating CD4^+^ T cells from NHB patients consistently exhibited enrichment of T cell activation (including α/μ T cell activation and CD28 co-stimulation), proliferation, differentiation, antigen receptor-mediated signaling, and cytokine-driven signaling (notably IL-2, IL-4/IL-13, and IL-6 pathways), along with chemotaxis and leukocyte trafficking across disease states (Fig. 6a-c). However, the balance between effector and regulatory programs differed by disease state. In the high-risk cohort, CD4^+^ T cells from NHB patients showed extensive activation, including Th1-associated cytokine signaling (IFN-γ, IL-12) and chemotaxis, yet simultaneous engagement of negative immune regulatory and tolerance pathways, indicating a stimulated but immunologically imbalanced state (Fig. 6a). In DCIS, CD4^+^ T cells from NHW patients displayed a more restricted profile, with enhanced activation and differentiation but relatively attenuated type I interferon signaling, cytotoxicity, ROS/RNS production, and tissue-specific effector responses, suggesting an antigen-responsive but comparatively controlled immune state (Fig. 6b). In breast cancer, CD4^+^ T cells from NHB patients demonstrated activation and cytokine responsiveness, but with prominent homeostatic and suppressive programs (including IL-10 signaling and multiple negative regulatory pathways) and reduced TNFR2 non-canonical NF-κB signaling, suggesting activation constrained by regulatory mechanisms (Fig. 6c). Flow cytometric analysis supported increased HLA-DR and PD-1 expression on naïve CD4^+^ T cells, and higher LAG-3 with lower CTLA-4 expressions across CD4^+^ T cell subsets in NHB patients (Supplementary Fig. S5a), consistent with heightened activation coupled with compensatory inhibitory regulation. Overall, these patterns suggest immune stimulation and dysregulation in the high-risk state, a relatively contained immune activation state in DCIS, and persistent activation but stronger immunoregulatory and homeostatic restraint in breast cancer in NHB relative to NHW patients.

**Figure 6.**
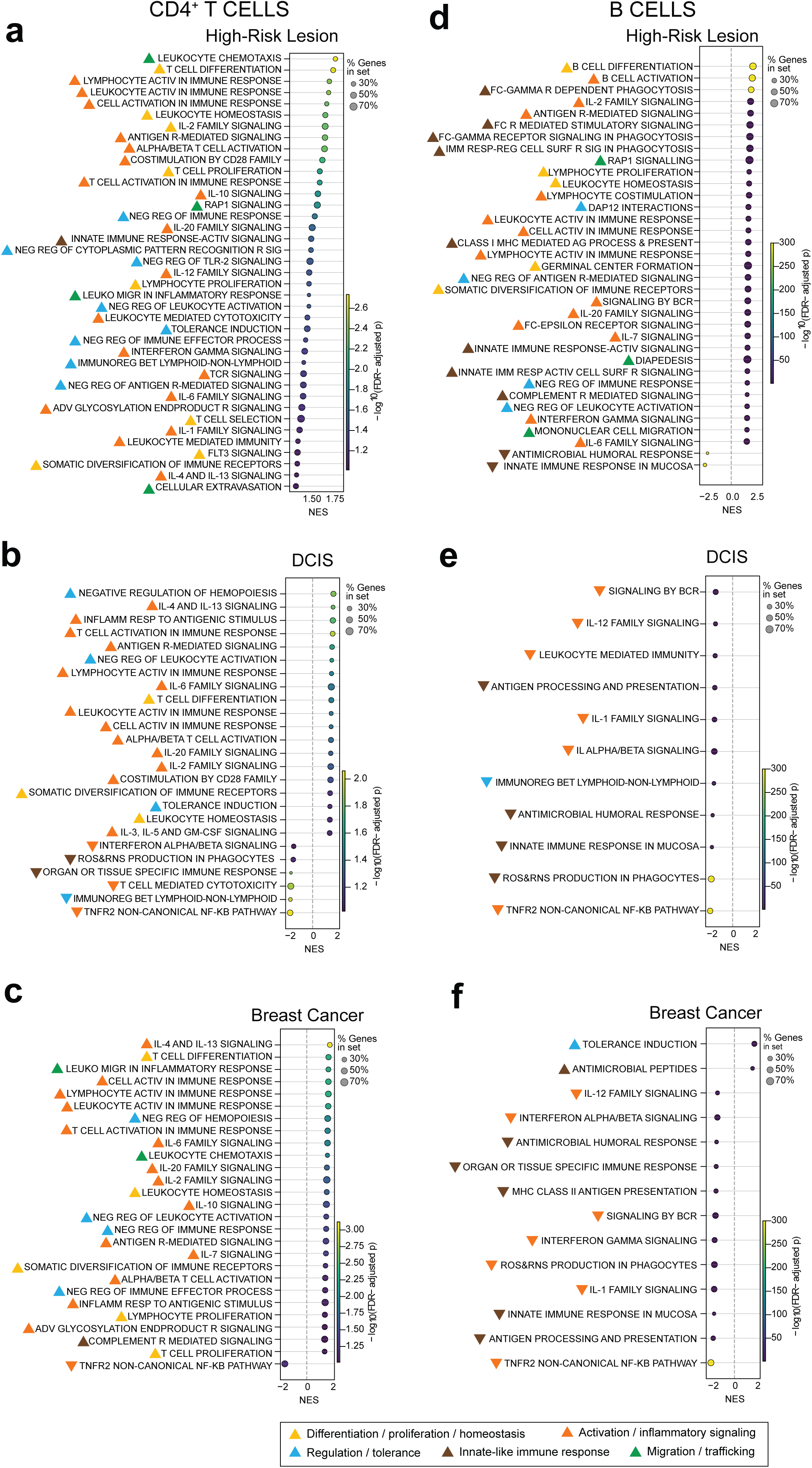
Race-associated pathways across disease states in CD4 + T cells and B cells. **a-f**, Pathways enriched (positive NES) and depleted (negative NES) in CD4^+^ T cells (a-c) and B cells (d-f) from NHB relative to NHW patients with HRL (a, d), DCIS (b, e), and BC (c,f) using GSEA and ImmPort databases. Bubble size, percentage of leading-edge genes (genes driving the enrichment signal) in the pathway; bubble color, -log10 FDR-adjusted P value; NES, normalized enrichment score. Colored arrows indicate pathways comprising general immune functional categories.

B cells from NHB relative to NHW patients exhibited distinct immune programs across disease states. In the HRL cohort, B cells from NHB patients demonstrated relative enrichment of activation, differentiation and proliferation, including B cell receptor (BCR) signaling, germinal center formation, somatic diversification, antigen processing and presentation, and Fc receptor-mediated signaling, along with cytokine pathways (IL-2, IL-6, IL-7, IFNγ) and migration-related processes (Fig. 6d). These results suggest a highly activated, antigen-experienced and immune complex-responsive state with concurrent regulatory feedback. However, in NHB patients with DCIS and invasive breast cancer, this B cell activation signature was largely absent, with downregulation of BCR signaling, antigen presentation, inflammatory cytokine pathways, and ROS/RNS production, which promotes BCR-signal amplification(24) (Fig. 6e,f), consistent with a comparatively quiescent or suppressed B cell compartment. Enrichment of tolerance induction and antimicrobial peptide programs in NHB patients with breast cancer further suggested a shift toward immune regulation and limited innate-like activity (Fig. 6f). Consistent with this finding, flow cytometry revealed reduced CCR7 and CXCR5 expression across B-cell subsets in NHB patients with breast cancer (Supplementary Fig. S5b), indicating altered homing and positioning.

Collectively, these analyses reveal that race-associated transcriptional differences in circulating myeloid (monocytes and DCs) and lymphoid (CD4^+^ T and B) cells are characterized by heightened immune activation coupled with impaired effector function and increased immunoregulatory signaling in NHB relative to NHW patients. These immune patterns in NHB patients are consistent with those observed in chronic inflammatory states.

### Population-level immune signature (IMM-POP) identifies an immunosuppressive immunotype in external datasets

We sought to generate a gene expression signature that captures transcriptional features unique to NHB versus NHW breast cancer patients. Starting with all race-associated DEGs identified in monocytes, dendritic cells, CD4^+^ T cells, and B cells from breast cancer patients (Fig. 4), we first aggregated gene expression across PBMCs from the same patients to generate pseudobulk RNA-seq data, and selected genes that remained significantly differentially expressed by race; these genes were defined as a population-level immune signature (IMM-POP; Fig. 7a). The derived IMM-POP comprised 37 genes with higher expression in NHB than NHW breast cancer patients (NHB-high IMM-POP genes) and 57 genes with lower expression in NHB than NHW breast cancer patients (NHB-low IMM-POP genes) (Fig. 7b, Supplementary Fig. S6a,b).

**Figure 7.**
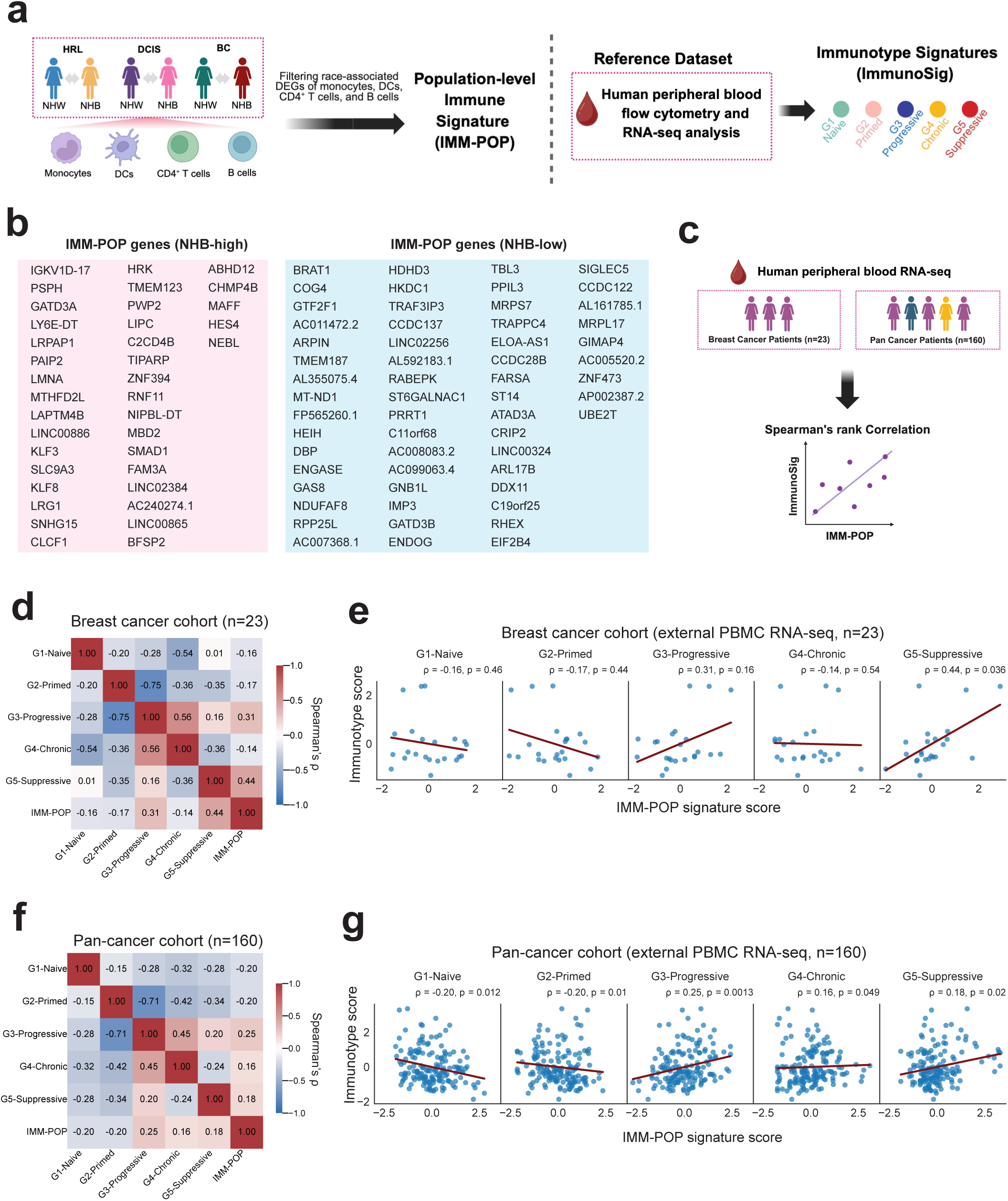
A population-level immune signature (IMM-POP) related to immunosuppression in NHB breast cancer patients. **a**, Workflow of IMM-POP signature generation and evaluating its relationship to published immunotype signatures(25): G1-naïve, G2-primed, G3-progressively activated, G4-chronically activated, and G5-suppressive. **b,** List of IMM-POP signature genes; 37 genes enriched and 57 genes reduced in NHB vs. NHW BC patients. **c**, Workflow for evaluating IMM-POP and immunotype signatures in two external datasets. **d-g**, Correlations between IMM-POP and G1–G5 immunotype signature scores in an external breast cancer cohort (n = 23 patients) (d, e) and a pan-cancer cohort (n = 160 patients) (f, g). Spearman (ρ) correlation coefficients (d, f) and original scores with fitted regression lines and Spearman’s ρ and p-values (e, g). BioRender. Chen, F. (2026) https://BioRender.com/8am5n3f and https://BioRender.com/ujnq4hq).

To understand whether IMM-POP relates to functional immune phenotypes (immunotypes), we quantified co-expression correlation of IMM-POP with known immunotype signatures (ImmunoSig) derived from a reference human PBMC multi-omics dataset(25) (Fig. 7a,c). ImmunoSig includes five signatures, each representing a distinct systemic immune status: naïve (G1; resting/immature), primed (G2; activated but not yet effector), progressively activated (G3; effector differentiation), chronically activated (G4; persistent stimulation and inflammation), and immunosuppressive (G5; regulatory and exhaustion-associated)(25) (Fig. 7a). To determine whether IMM-POP relates to any of the 5 ImmunoSigs, we tested their co-expression patterns using two PBMC bulk RNA-seq datasets, a breast cancer cohort consisting of newly diagnosed, pre-treatment breast cancer patients (n = 23), and a pan-cancer cohort consisting of patients with multiple cancer types (n = 160)(25) (Fig. 7c).

In the breast cancer cohort, IMM-POP scores positively correlated with the G5-immunosuppressive immunotype and had no significant associations with the other four immunotypes (Fig. 7d,e). In the pan-cancer cohort, IMM-POP positively associated with the G3-progressively activated, G4-chronically activated, and G5-immunosuppressive immunotypes, and negatively correlated with the G1-naïve and G2-primed immunotypes (Fig. 7f,g). Collectively, these findings suggest that transcriptional features distinguishing NHB from NHW breast cancer patients are linked to a chronically activated and immunosuppressive/exhaustion-like immune state.

## Discussion

This study provides a peripheral immune atlas from racially balanced cohorts spanning precancerous and invasive breast cancer, revealing marked transcriptional differences between populations across disease states and underscoring the close relationship between systemic immunity and cancer development. In this study, race-associated transcriptional differences in systemic immunity were detectable even in the high-risk stage and became more pronounced and biologically distinct in invasive breast cancer, suggesting that immune differences arise early and may be further shaped during disease evolution. The ability to distinguish early lesions from invasive breast cancer using PBMC profiling supports the potential utility of race-appropriate peripheral blood-based biomarkers for monitoring breast cancer progression, particularly in settings where tissue is limited or unavailable.

Race-associated transcriptional differences were most pronounced in monocytes, DCs, CD4^+^ T cells, and B cells, indicating that immune lineage composition and function are key domains of biological variation across groups. Across these four immune lineages, enrichment patterns differed by disease state, with the breast cancer cohort in particular, showing a concurrent enrichment of inflammatory and immune regulatory programs in NHB relative to NHW patients. This combined activation-regulation signature is consistent with a state of chronic immune stimulation accompanied by compensatory immune suppression. This interpretation is further supported by the immunotype analyses from independent cohorts, where higher IMM-POP scores in breast cancer were selectively associated with an immunosuppressive signature, and in pan-cancer, which showed positive associations with progressively activated, chronically activated, and immunosuppressive immunotypes. These observations are consistent with a state of chronic inflammation(5) and prior reports of race-associated immune differences in other disease contexts, including malaria(26), systemic lupus erythematous(27), and lung cancer(28), in which Black individuals similarly exhibit heightened inflammatory responses relative to White individuals.

Chronic inflammation (inflammaging) is a defining feature of immune aging(7,29,30). Consistent with this, NHB patients across disease states exhibited higher proportions of non-classical monocytes relative to NHW patients, a classic feature of immune aging(7). These observations are reminiscent of the weathering hypothesis(31), which posits that in the United States, Black individuals experience accelerated aging and enhanced allostatic load compared with White individuals. Our data suggest an “immune weathering” that manifests in NHB women at earlier breast disease states than NHW women. Future studies are therefore warranted to investigate whether early immune weathering contributes to the younger onset of breast cancer experienced by NHB individuals(2). In this context, the IMM-POP signature, which reflects features of immune aging, may serve as a useful metric for patient stratification in studies of early immune aging and cancer development.

This study also has limitations. First, it is cross-sectional rather than longitudinal; hence, observed transcriptional differences across disease states may be impacted by patient heterogeneity. Second, the DCIS cohort was relatively small (4 NHB and 6 NHW patients), which may limit statistical power for race-stratified comparisons of immune cell composition. Finally, longitudinal clinical follow-up, including recurrence and survival data, will enable future analyses to determine whether baseline peripheral immune profiles can be used to predict clinical outcomes and improve risk stratification across racially diverse patient groups. Such efforts will ultimately improve prevention, treatment, and outcomes for all populations suffering from breast cancer.

## Methods

### Cohort recruitment and clinical information collection

The cohort consisted of pre-treatment NHW and NHB patients diagnosed with high-risk breast lesions (HRL), DCIS, or breast cancer. The HRL patients were recruited from the Breast Cancer Personalized Risk Assessment, Education and Prevention (B-PREP) program (Dana-Farber/Harvard Cancer Center protocol IRB #13-325)(9), where HRL status was defined by the presence or history of a biopsy-proven atypical ductal hyperplasia, atypical lobular hyperplasia, or lobular carcinoma in situ. The DCIS and breast cancer patients were enrolled through Project SHARE (Dana-Farber/Harvard Cancer Center protocol IRB #93-085). The HRL, DCIS, and breast cancer patients were designed to be age-matched by race and disease state. Deidentified demographic and clinical information was retrieved from electronic medical records on EPIC, including age at blood collection, diabetes (present or past medical history of diabetes), and obesity status (present or past medical history of obesity and/or BMI ≥ 30 kg/m^2^). Additional clinical and pathological information was obtained from the electronic medical records and pathology reports for breast cancer or DCIS patients, which included TNM categories, tumor grade, estrogen receptor (ER) status, progesterone receptor (PR) status, and human epidermal growth factor receptor 2 (HER2) status. The use of human samples was approved by the Dana-Farber/Harvard Cancer Center Institutional Review Board. All participants provided written informed consent before sample collection for blood use for research purposes. The conduct of this research adhered to institutional and/or national ethical guidelines and the principles of the 1964 Helsinki Declaration and its subsequent amendments.

### Single-cell RNA-seq library preparation and sequencing

Freshly collected PBMC aliquots were slow frozen overnight at -80 °C, and cryopreserved in liquid nitrogen until needed for analysis. At time of sequencing, the cells were washed and resuspended in phosphate-buffered saline with 0.04% bovine serum albumin at a concentration of 1000 cells/µL, and approximately 26,000 cells were loaded onto a 10x Genomics Chromium^TM^ X instrument (10x Genomics) according to the manufacturer’s instructions. The scRNA-seq libraries were processed using Chromium Next GEM Single Cell 5’ HT Kit v2 (10x Genomics, PN-1000356). Quality controls for amplified cDNA libraries and final sequencing libraries were performed using Bioanalyzer High Sensitivity DNA Kit (Agilent, Cat#: 50674626). The sequencing libraries for scRNA-seq were normalized to 4nM concentration and pooled. The pooled sequencing libraries were sequenced on the Illumina NovaSeq6000 S4 300 cycle platform. The sequencing parameters were: Read 1 of 28bp, Read 2 of 90bp, Index 1 of 10bp and Index 2 of 10bp.

### scRNA-seq data preprocessing

Single cell RNA-sequencing was performed at the DFCI Translational Immunogenomics Lab (TIGL). The sequencing data were demultiplexed, and the scRNA-seq data were aligned to GRCh38-2020-A reference using cellranger v7.1.0 (10x Genomics). We used Scrublet (v0.2.3) to detect doublets(32). We used Azimuth (0.5.0) with PBMC reference(11) to annotate cell types and to compute expression of surface protein markers of major immune lineages. For each sample, we excluded low quality cells (expressing ≤1000 genes, or with mitochondrial percentage ≥10%) and lowly-expressed genes (expressed by ≤3 cells). We excluded cells predicted as doublet by Scrublet, or those which had normalized expression of erythrocyte marker genes (*HBB*, *HBD*) or platelet marker genes (*PF4*, *SDPR*, *GNG11*, *PPBP*) of e^2^ – 1 or higher(11). We also excluded cells with main cell type prediction score < 0.9 (level 1 cell type prediction score from Azimuth), and cells assigned as hematopoietic stem cell, progenitor cell, platelet, or innate lymphoid cells from Azimuth annotation (Supplementary Fig. S1a–d). We then followed standard workflow of Seurat (v5.3.1)(33) for normalization, finding variable genes, scaling, dimensional reduction by principal component analysis (PCA) and Uniform Manifold Approximation (UMAP), data integration, and Louvain clustering. One patient was excluded after the cell type annotation step from the original cohort due to an unusually high proportion of mature B cells (approximately 30% of total PBMC) suggestive of unidentified hematologic conditions.

### Cell type annotation and marker gene identification from scRNA-seq data

Major cell types were assigned using level 1 cell type predictions from Azimuth (B cells, CD4^+^ T cells, CD8^+^ T cells, other T cells, dendritic cells, monocytes, and NK cells). Within each major cell type, we repeated the Seurat workflow as described and performed Louvain clustering with resolution ranging from 0.5 to 1. Highly expressed genes of each cluster (cluster marker genes) were identified using Seurat’s *FindAllMarkers* function, and were used to annotate immune cell subtypes of the corresponding cluster and thereby to determine the final clustering and cell type annotations, using subtype classification from literature.

### Cell Proportion analysis

Peripheral blood cell proportions were calculated for each immune cell subtype within the corresponding lineage (B cells, T cells, dendritic cells, monocyte, and NK cells). We used Mann-Whitney U test to compare cell type percentage of major cell type between patient groups.

### Differential gene expression and gene set enrichment analysis by disease state and by race

We first calculated normalized expression for each patient and each cell type using *AggregateExpression* function from Seurat to generate the *pseudobulk* data. Then, we applied edgeR (v4.4.2)(34) to the pseudobulk data to identify differentially expressed genes (DEGs) between patient groups, using a design formula including an interaction term between race and disease state, adjusting for age, diabetes, and obesity status. Significant DEGs were defined by FDR-adjusted p-value < 0.1 from edgeR result. For pathway enrichment analysis, we used fgsea package (v1.32.4)(35) based on gene ranking by signed –log_10_p-values (positive if upregulated, negative if downregulated) from the edgeR differential expression analysis, and significant pathways were defined by FDR < 0.05 from the fgsea results.

### Co-varying neighborhood analysis and pathway scoring

We performed co-varying neighborhood analysis (CNA v0.2.2)(22) in scRNA-seq data of each major immune cell type (B cells, CD4^+^ T cells, CD8^+^ T cells, other T cells, dendritic cells, monocytes, and NK cells) for each patient feature of interest, including age, race, disease state, diabetes, and obesity status. Within the main cell types showing significant association with race (the monocytes, dendritic cells, CD4^+^ T cells, and B cells), we also applied the same CNA analysis in each disease state (HRL, DCIS, and breast cancer) in each major immune cell lineage respectively. Significance of associations was defined according to original publication of CNA(22), namely, p < 0.05 for significant global association, and FDR < 0.1 for significant cell-level (local) associations. For pathway enrichment stemming from CNA analysis, we used the gseapy (v1.1.9) package(36) to conduct GSEA analysis using gene ranking defined by Spearman correlation coefficient between normalized gene expression and the cell-level association coefficient with race (NHB vs. NHW patients). Significant pathways were selected based on an FDR-adjusted P value < 0.1 in GSEA and further filtered to retain only those relevant to the immune cell lineage analyzed.

### Developing the population-level immune signature (IMM-POP)

To derive the population-level immune signature (IMM-POP), we first identified the differentially expressed genes (FDR<0.05) from monocytes, dendritic cells, CD4^+^ T cells, and B cells in NHB compared to NHW breast cancer patients (Seurat). Then we aggregated gene expression from all PBMCs for each patient (*AggregateExpression* function from Seurat), and selected genes which had higher or lower expression in NHB than NHW breast cancer patients (NHB-high IMM-POP genes and NHB-low IMM-POP genes respectively) (Mann–Whitney U test, FDR < 0.05). Expression of each gene from the IMM-POP was compared between patient groups using the Mann–Whitney U test.

### Expression of IMM-POP and immunotype signatures in external datasets

Two public PBMC bulk RNA-seq datasets were used: dataset 1 (n=23) was from a cohort of newly-diagnosed, pre-treatment breast cancer patients(37) and dataset 2 (n=160) consisted of patients with multiple cancer types(25). The definition of immunotype signatures (G1-naïve, G2-primed, G3-progressively activated, G4-chronically activated, and G5-immunosuppressive) and immunotype signature scores of these two datasets were previously published(25). For each sample, we used GSVA package (single-sample GSEA mode) (v2.0.7)(38) to calculate signature scores for both IMM-POP and immunotype signatures. Specifically, we calculated an IMM-POP score for each patient by subtracting the ssGSEA score of NHB-low IMM-POP genes from that of NHB-high IMM-POP genes, and immunotype scores as ssGSEA scores for the G1–G5 signatures. Spearman correlation coefficient and p values were computed between scores of each immunotype signature and the IMM-POP signature across patients.

### Spectral flow cytometry cohort analysis

The spectral flow cytometry cohort, independent of the scRNA-seq cohort, consisted of pre-treatment early-stage ER+/HER2- breast cancer patients enrolled in the DAPHNe trial (NCT03716180) and the MARGOT trial (NCT04425018). Antibody panel for detecting T/B cell markers was used. The final dataset included 18 NHB and 26 NHW patient samples, including 13 NHB and 13 NHW patients overlapped with the scRNA-seq cohort. Detailed experimental and analysis steps of the spectral flow cytometry were described in previous publication(10). Briefly, cryopreserved freshly collected PBMCs were thawed, stained with fluorochrome-labeled antibodies from the T/B panel, and flow cytometry was performed. Data was preprocessed with Cytek Aurora SpectroFlo software (v2.2) for unmixing, with OMIQ (app.omiq.ai) for quality control, filtering, and gating. Arcsinh scaled mean fluorescence intensity (MFI) data was exported from OMIQ under default settings (arcsinh scaling) for each sample and gated cell population. Cell subtypes were annotated based on sequential gating via specific protein marker MFI values in the published workflow(39).

### Statistical analysis

All statistical tests were two-sided. Adjustment for multiple comparisons was performed using the Benjamini–Hochberg false discovery rate (FDR) method. Statistical analyses were conducted in Python (v3.12) and R (v4.4) using the packages described above.

## Data Availability

The raw scRNA-seq data (fastq) files were deposited at the Sequence Read Archive (SRA) with BioProject accession PRJNA1280085. Processed scRNA-seq data was deposited at the Gene Expression Omnibus (GEO) with accession GSE318467. The raw and processed flow cytometry data was deposited at zenodo at 10.5281/zenodo.18509855.

## Code Availability

Code for data analysis and plotting was deposited on GitHub (https://github.com/ChelseaCHENX/Immune-Weathering-scRNA-seq).

## Authors’ Contributions

E.A.M. and S.S.M. designed the study. E.R.O. and A.R. obtained and processed the blood samples. R.T. supervised sample annotation and storage. E.R.O. and M.S. prepared samples for RNA sequencing, performed flow cytometry, and preprocessed the data. F.C. analyzed and interpreted the data. A.M.P. and P.vG. supervised data analysis and interpretation. F.C. wrote the manuscript and all authors edited the paper. E.A.M. and S.S.M. provided funding. All authors read, edited, and approved the paper.

## Acknowledgments

E.A.M. is supported by the Harvard Ludwig Award, the Komen Award (SAC210204), and the Parker Institute for Cancer Immunotherapy Award (C-03159). S.S.M. is supported by the Victoria’s Secret Global Fund for Women’s Cancers Rising Innovator Research Grant in partnership with Pelotonia and the AACR (23-30-73-MCAL), Samuel Waxman Cancer Research Foundation with the Mark Foundation for Cancer Research. We thank members of the van Galen, McAllister, and Mittendorf laboratories for helpful discussions and input, and the DFCI Translational Immunogenomics Laboratory for technical guidance. We are grateful to the patient donors and their families.

## Supplementary Figure Legends

**Supplementary Fig. S1 Quality control metrics of scRNA-seq data and expression patterns of marker proteins and marker genes of PBMC cell types. a-d**, Quality control metrics of PBMC scRNA-seq, including per-cell gene counts (a), unique molecular identifier counts (b), mitochondrial gene percentage (c), and major cell type prediction scores (d). **e,** Imputed expression of protein markers of major immune cell types. f. Expression of marker genes of each functional minor PBMC cell type.

**Supplementary Fig. S2 PBMC composition across disease states and racial groups. a-b,** Percentage of major immune cell types within total PBMCs for comparisons in Figure 2a (a) and Figure 2j (b). **c-d,** Percentage of PBMC cell types within the indicated parental immune lineage for comparisons in Figure 2a (c) and Figure 2j (d). All cell types with no significant differences in comparisons by Mann-Whitney U test were shown. HRL, high-risk breast lesion; DCIS, ductal carcinoma in-situ; BC, breast cancer.

**Supplementary Fig. S3 Transcriptional differences in peripheral blood immune cells across disease states in each racial group**. **a-b,** Gene set enrichment analysis of the Molecular Signatures Database (MSigDB) hallmark pathways in each cell type from comparisons between disease states (DCIS vs. HRL, BC vs. DCIS, and BC vs. HRL) in NHW (a) and NHB patients (b). MSigDB Hallmark pathways not displayed in Figure. 3 were shown. Box color represents normalized enrichment score (NES); asterisks indicate GSEA FDR-adjusted p value < 0.05.

**Supplementary Fig. S4 Subtype compositions of immune cell transcriptional states that associate with race**. **a-d,** UMAP plots of the single-cell transcriptomic data of monocytes (a), dendritic cells (b), CD4^+^ T cells (c), and B cells (d), colored by cell subtypes. **e-h,** Cell subtype compositions of immune cells having a transcriptional state associated with race: positive association with NHB / enriched in NHB (left) and negative association with NHB / depleted in NHB (right).

**Supplementary Fig. S5 Protein expressions by race in CD4^+^ T cells and B cells from breast cancer patients. a-b,** Mean fluorescence intensity (MFI) values of marker proteins in CD4^+^ T cells (a) and B cells (b) from full-spectrum flow cytometry data of breast cancer patients. Mann-Whitney U test; * p ≤ 0.05, ** p ≤ 0.01, *** p ≤ 0.001, **** p ≤ 0.0001, ns p > 0.05. CD4^+^ TCM, central memory CD4^+^ T cells; CD4^+^ TEM, effector memory CD4^+^ T cells; CD4^+^ CTL, cytotoxic CD4^+^ T cells; Treg, regulatory T cells.

**Supplementary Fig. S6 Expression of the IMM-POP signature by race in breast cancer patients.** a-b, Expression of the 37 IMM-POP genes enriched (a) and 57 IMM-POP genes reduced (b) in NHB vs. NHW BC patients. Expression presented on patient level by pseudobulk data. Mann-Whitney U test; * p ≤ 0.05, ** p ≤ 0.01, *** p ≤ 0.001, **** p ≤ 0.0001, ns p > 0.05.

