## Supplementary figures and images for "A Single-Cell Peripheral Immune Atlas Spanning High-Risk Lesions to Invasive Breast Cancer in Black and White Women"

### Supplementary Fig. S1

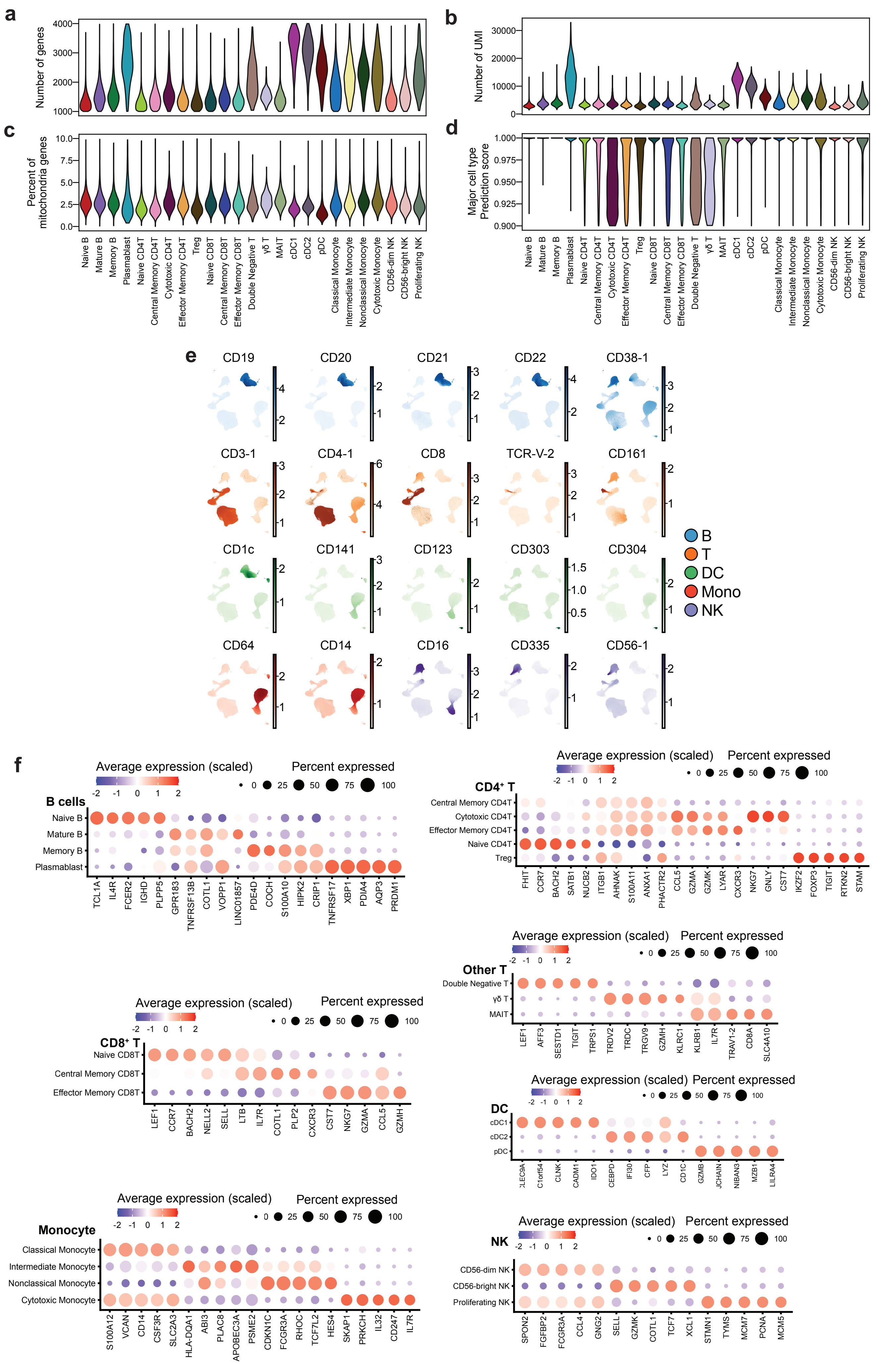

### Supplementary Fig. S2

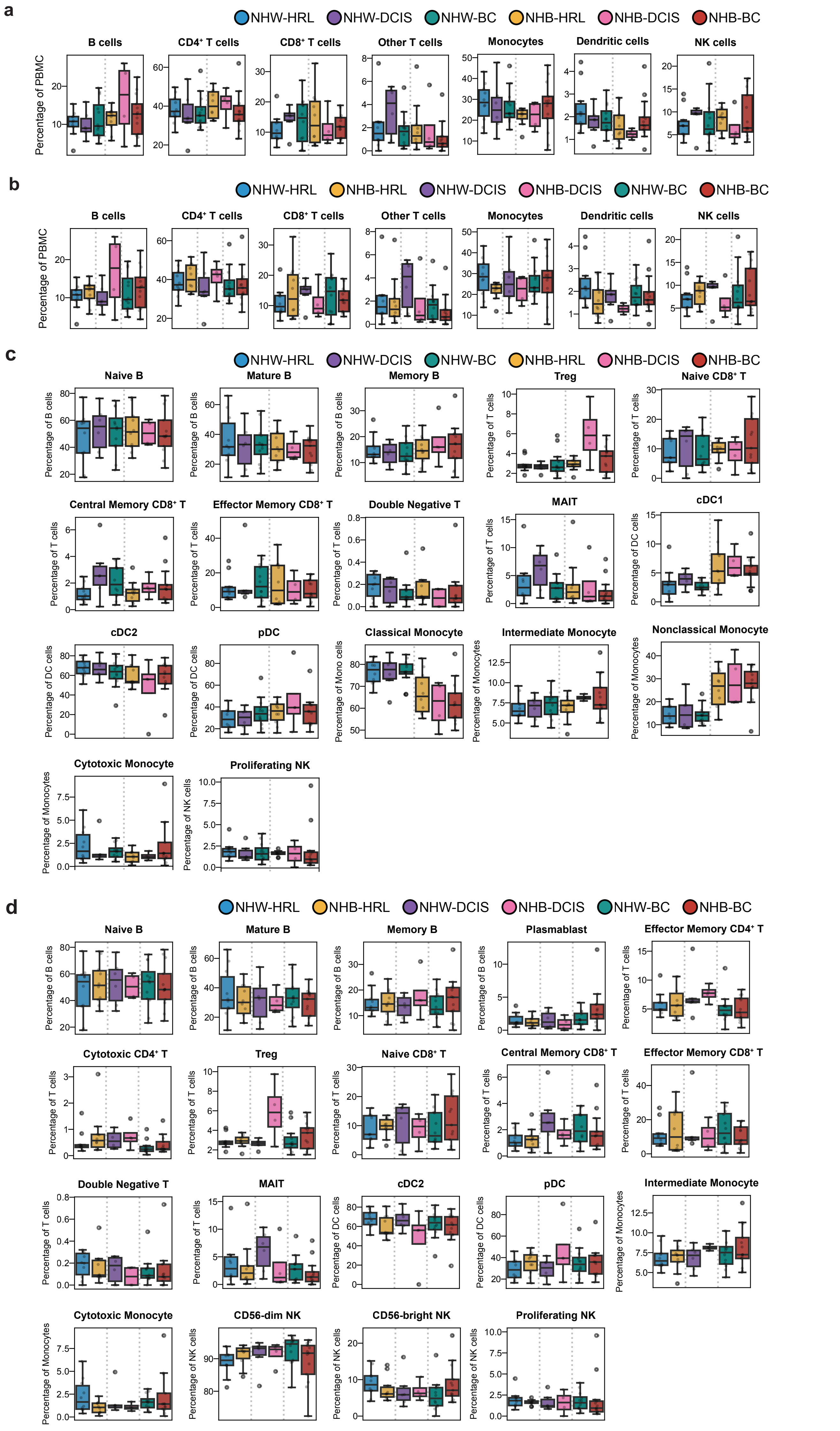

### Supplementary Fig. S3

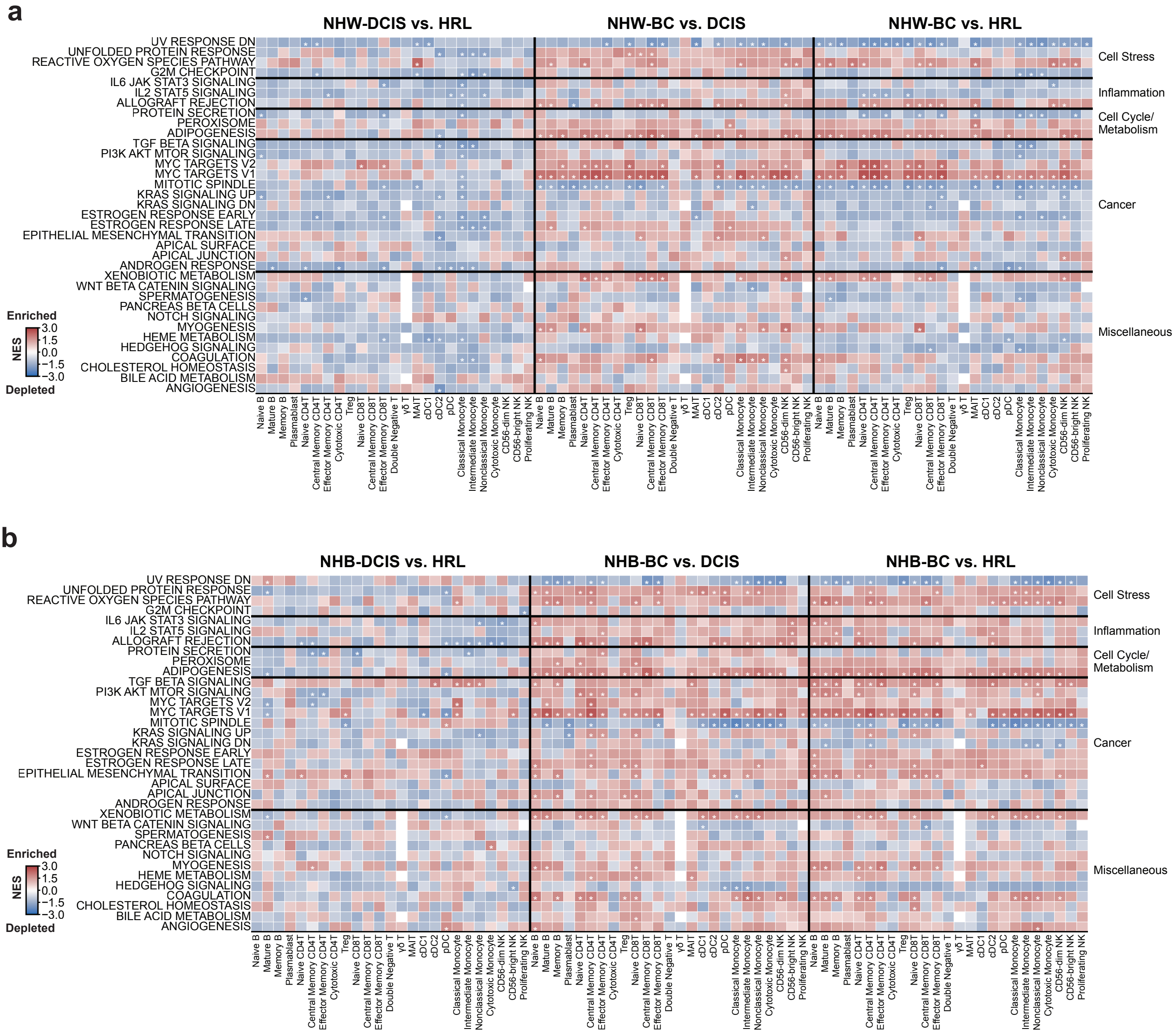

### Supplementary Fig. S4

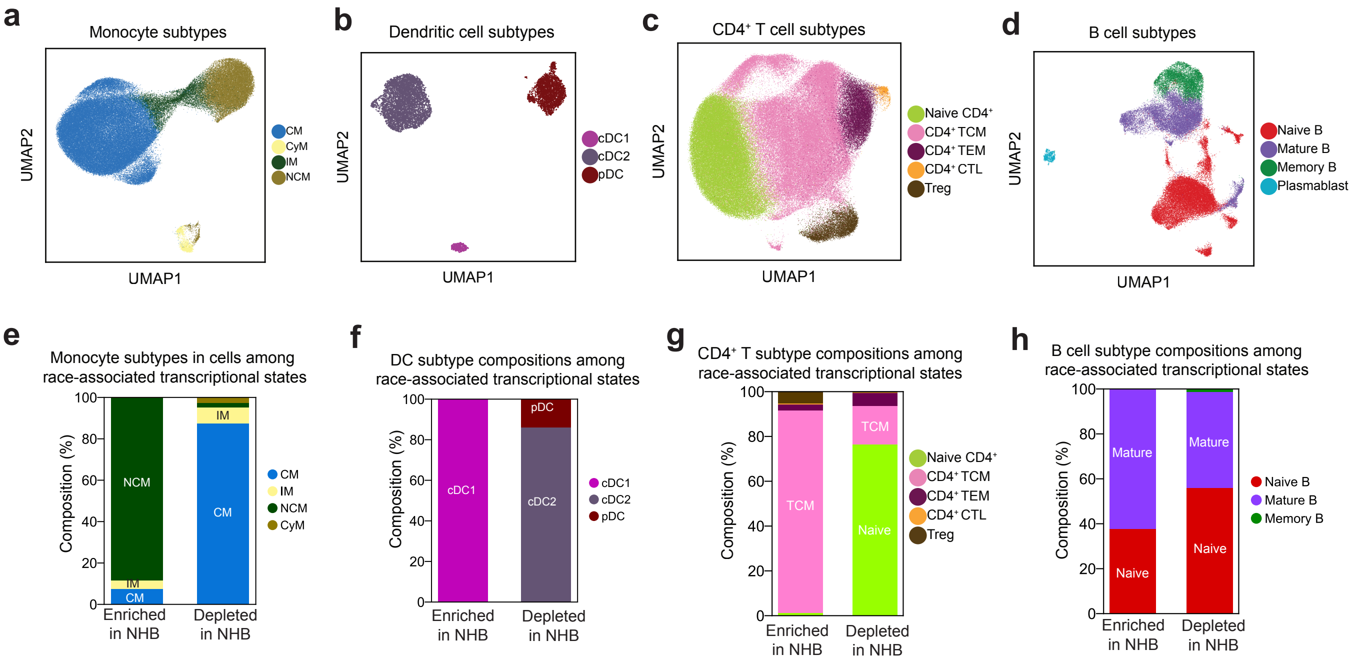

### Supplementary Fig. S5

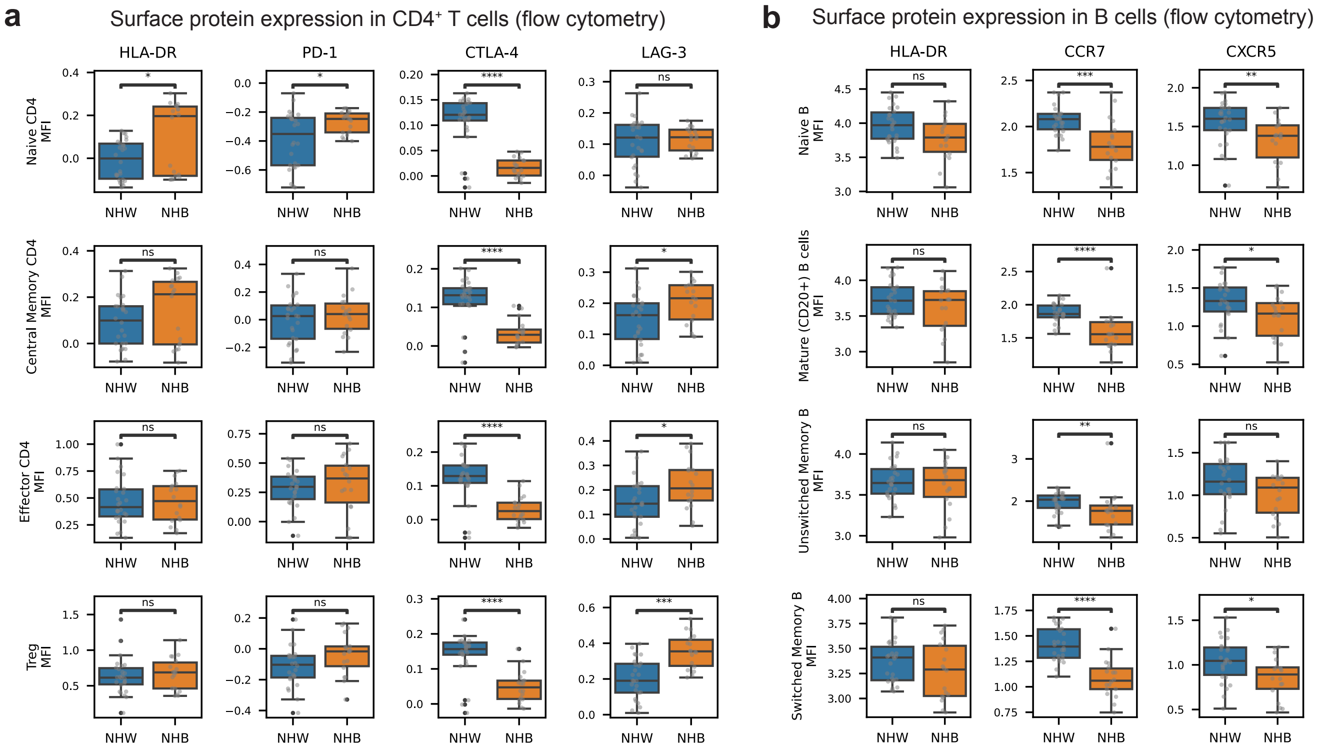
