## Supplementary Fig. S6 for "A Single-Cell Peripheral Immune Atlas Spanning High-Risk Lesions to Invasive Breast Cancer in Black and White Women"

### ExDat Fig. 6

**a**

#### Patient-level expression of NHB-high IMM-POP genes by race

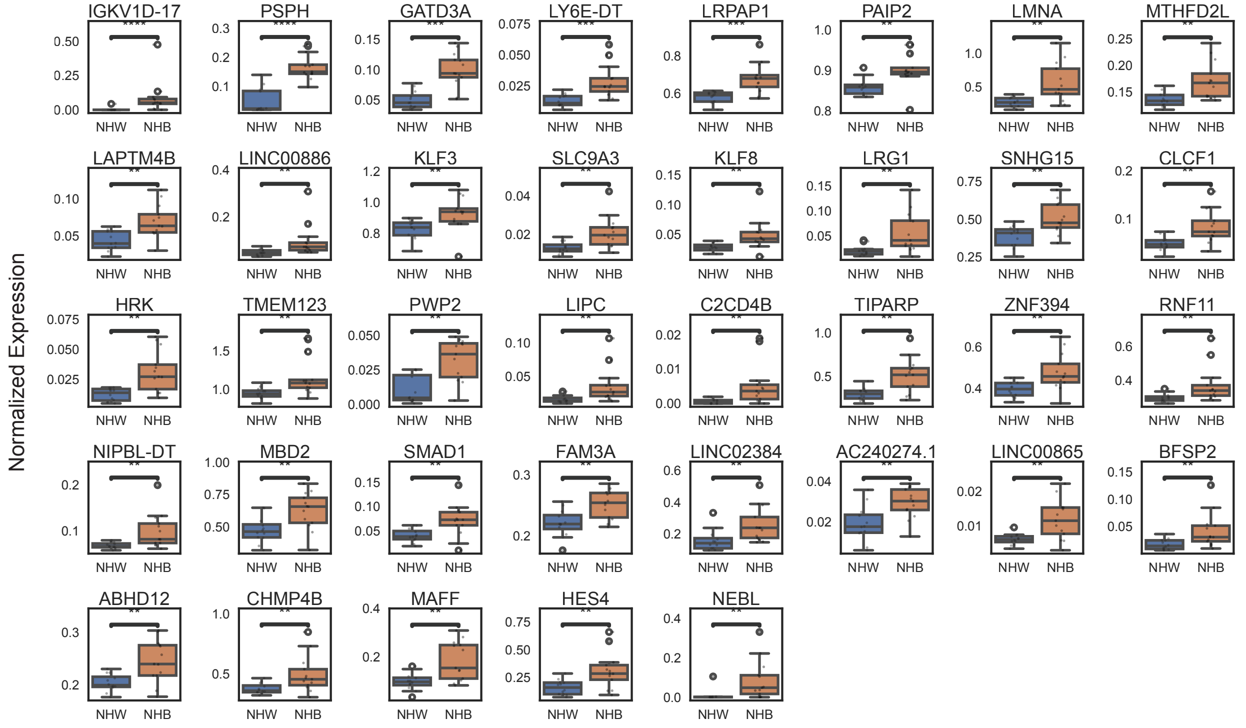


**b**

#### Patient-level expression of NHB-low IMM-POP genes by race

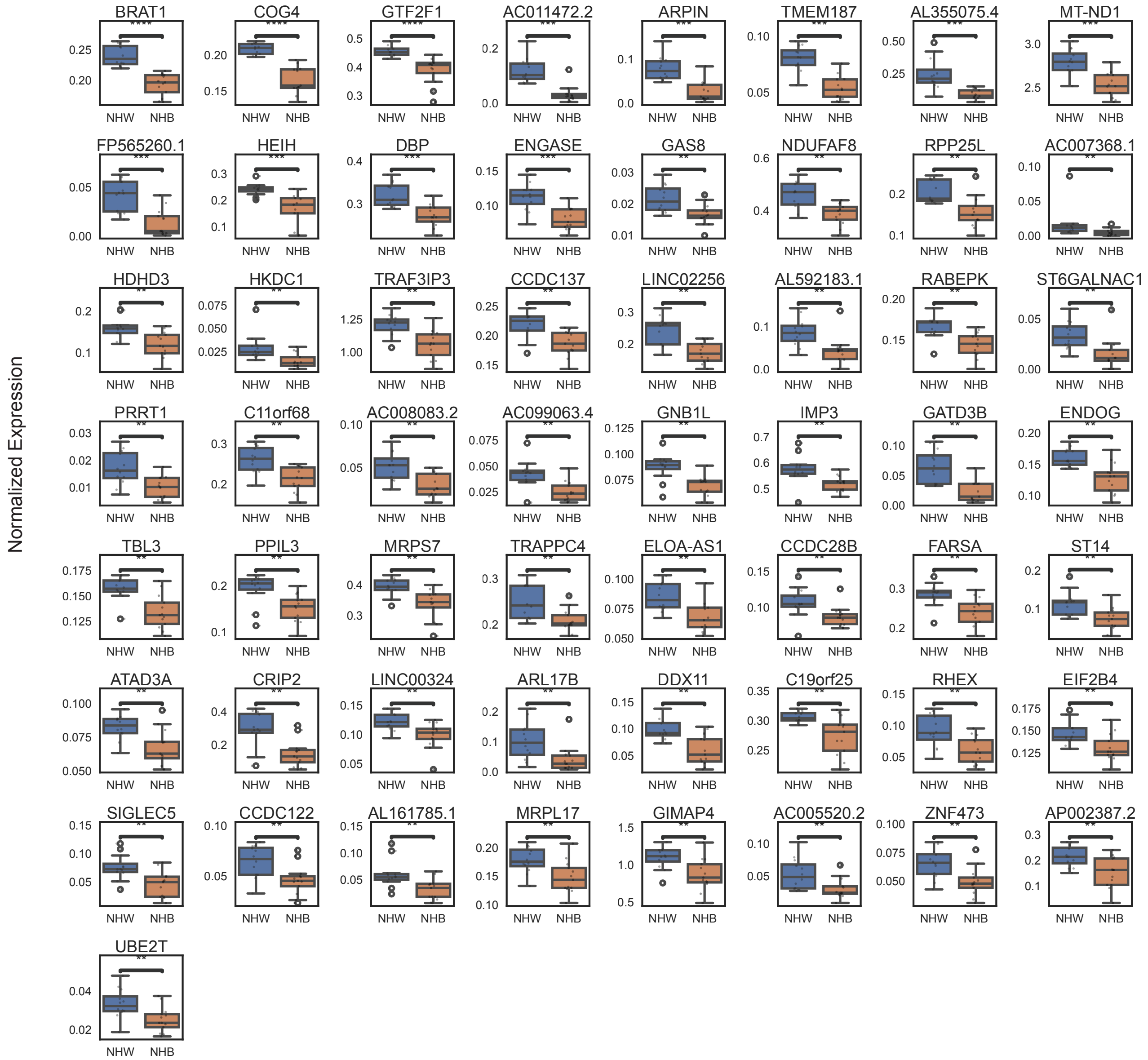
